# An optimized high-yield protocol for expression and purification of monomeric full-length BAX protein

**DOI:** 10.1101/2023.10.16.562589

**Authors:** Yiyang Chen, Jesse D. Gelles, Jarvier N. Mohammed, Jerry Edward Chipuk

## Abstract

Diverse developmental signals and pro-death stresses converge on regulation of the mitochondrial pathway of apoptosis. BAX, a pro-apoptotic BCL-2 effector, directly forms proteolipid pores in the outer mitochondrial member to activate the mitochondrial pathway of apoptosis. BAX is a viable pharmacological target for various human diseases, and increasing efforts have been made to study the molecular regulation of BAX and identify small molecules selectively targeting BAX. However, generating large quantities of monomeric and functionally-competent BAX has been challenging due to its aggregation-prone nature. Additionally, there is a lack of detailed and instructional protocols available for investigators who are not already familiar with recombinant BAX production. Here, we present a comprehensive high-yield protocol for expressing, purifying, and storing functional recombinant BAX protein. We utilize an intein-tagged BAX construct and employ a two-step chromatography strategy to capture and purify BAX, and provide example standard assays to observe BAX activation. We also highlight best practices for handling and storing BAX to effectively preserve its quality, shelf-life, and function.

## Introduction

Apoptosis is a fundamental physiological process essential for tissue development and homeostasis. The BCL-2 family of proteins governs the mitochondrial pathway of apoptosis by regulating mitochondrial outer membrane permeabilization (MOMP), an event signifying as the “point of no return” and a cellular commitment to apoptotic death. Following MOMP, several apoptogenic factors are released from the mitochondrial inter-membrane space into the cytosol, such as cytochrome *c*, which triggers the caspase cascade for cellular dismantling.

BAX, a proapoptotic BCL-2 effector protein, directly mediates MOMP by forming proteolipid pores on the outer mitochondrial membrane (OMM). Nascent BAX exists as a soluble monomer, with its ɑ9 helix occupying the BH3-and C-terminal (BC) groove (Suzuki et al., 2000). In response to apoptotic stimuli, BAX is activated by a subset of BCL-2 family proteins (the “direct activators”, such as BIM) (Kuwana et al., 2002, Gavathiotis et al., 2008). Upon activation, BAX undergoes a series of conformational changes, including mobilization of the ɑ9 helix from the BC groove, which facilitates the translocation of BAX to mitochondria, oligomerization, and subsequent pore formation in the OMM (Antonsson et al., 2000, Gross et al., 1998). Given the central role of BAX in initiating apoptosis, there has been increasing efforts to study the structure-function relationship and molecular regulation of BAX, which has led to the discovery of several regulatory sites (Gavathiotis et al., 2008, Czabotar et al., 2013, Barclay et al., 2015). Furthermore, several studies have identified or developed small molecules capable of directly targeting BAX and modulating its function (Gavathiotis et al., 2012, Reyna et al., 2017, Pritz et al., 2017, Niu et al., 2017, Garner et al., 2019).

Several multi-domain BCL-2 family proteins are membrane-associated, thereby introducing technical complications for generation of their full-length recombinant protein. By contrast, BAX uniquely exists as a soluble, cytosolic protein in its inactive state, simplifying the process of generating recombinant protein and representing a biologically-relevant state for structure-function studies. Various biophysical and biochemical techniques commonly utilized in BAX research (e.g., NMR, ITC, fluorescence polarization, membrane permeabilization assays) require large quantities of highly-pure and functionally-competent BAX protein. Therefore, reliably generating high-quantity and high-quality recombinant BAX is indispensable for BAX research. However, due to the aggregation-prone nature of BAX, generating, handling, and storing large quantities of recombinant BAX is a challenge.

Here, we present a high-yield protocol for the generation of a high purity, monomeric, full-length recombinant human BAX. We utilize an intein-tagged BAX construct and employ a two-step chromatography strategy to capture and purify monomeric BAX. To validate the function of recombinant BAX, we conduct a series of gold standard assays to ensure BAX is highly responsive to various BAX activators. Furthermore, we optimized the storage condition to effectively preserve the high quality of BAX, ensuring consistency across a spectrum of experiments. Collectively, this comprehensive protocol is beneficial for investigators who are not familiar with recombinant BAX production and serves a useful resource for experienced investigators to compare best practices.

## Materials and Equipment

### Laboratory Equipment

- High-speed Centrifuge (e.g., Avanti J-E Series; Beckman coulter)
- Incubator shaker capable of holding 2 L flask (e.g., Innova44; New Brunswick Scientific)
- Liquid chromatography systems (e.g., ÄKTA pure™ 25 L1; Cytiva)
- Microfuge tube centrifuge (e.g., Legend micro 21R; Thermo Scientific)
- Microplate reader capable of absorbance and fluorescence (e.g., Synergy H1; BioTek)
- Probe Sonicator (e.g., Dismembrator 505; Fisher Scientific)
- Protein electrophoresis & western blot sets (e.g., Criterion Vertical Electrophoresis Cell; BioRad)
- Spectrophotometer capable of cuvette reading (e.g., Ultraspec7000; GE)
- Size exclusion column (e.g., HiLoad 16/600 Superdex 200 pg column; Cytiva)
- Table-top swinging bucket centrifuge with 50 ml tubes adapters (e.g., Legend XTR centrifuge; Thermo Scientific)
- Thermomixer (e.g., ThermoMixer; Eppendorf)

### Consumables and Labware

- 1.5 ml Eppendorf tubes (Cat. No. 4036-3204, USA Scientific)
- 1.5 ml semi-micro cuvettes (Cat. No. 14955127, Thermo Scientific)
- 2 L culture flask (Cat. No. CLS431256, Sigma)
- 5 ml syringe with luer-lock tip (Cat. No. 309646, BD)
- 50 ml conical centrifuge tubes (Cat. No. SC-200251, Santa Cruz)
- 96-well plate, flat bottom, black polystyrene (Cat. No. 3915, Corning)
- 96-well clear plates
- Amicon^®^ Ultra-4 centrifugal Filters, 10 kDa MWCO (Cat. No. UFC801024, Sigma Aldrich)
- Gravity chromatography column (25 mm × 200 mm) (Cat. No. SC-205557, Santa Cruz)
- Oak Ridge centrifuge tube (Cat. No. 3119-0050, Thermo Scientific)
- Petri dish, 100 × 15 mm (Fisher Scientific, catalog number: FB0875712)
- Serological pipettes of various size (Globe Scientific)

### Reagents

- Carbenicillin (Cat. No. 20871, Cayman Chemical)
- Chitin Resin (Cat. No. S6651L, New England Biolabs)
- Dithiothreitol (DTT) (Cat. No. DTT10, GoldBio)
- BL21 Star (DE3) cells (Cat. No. C601003, Thermo Fisher)
- GelCode™ Blue Stain Reagent (Cat. No. 24592, Thermo Scientific)
- Gel filtration molecular weight standard (Cat. No. 1511901, Bio-Rad)
- Glycerol (Cat. No. BP229-1, Fisher Scientific)
- 4-(2-hydroxyethyl)-1-piperazineethanesulfonic acid (HEPES) (Cat. No. A14777.30, Thermo Scientific)
- Isopropyl-β-D-thiogalactoside (IPTG) (Cat. No. I2481C, Gold Biotechnology)
- Luria-Bertani (LB) Agar, Miller (Cat. No. DF0445-17-4, Fisher Scientific)
- LB both (Miller’s) (Cat. No. BD 244610, Fisher Scientific)
- 1× Phosphate-buffered saline (PBS) [pH 7.4] (Cat. No. 10010-023, Gibco)
- Pierce™ protease inhibitor cocktail (Cat. No. A32965, Thermo Scientific)
- Potassium phosphate dibasic (K_2_HPO_4_) (Cat. No. BP363-500, Fisher Scientific)
- Sodium Chloride (NaCl) (Cat. No. 447300050, Thermo Scientific)
- Sodium phosphate monobasic (NaH_2_PO_4_) (Cat. No. BP329-1, Fisher Scientific)
- Terrific Broth (TB) Medium (Cat. No. DF0438-17, Fisher Scientific)

### Buffer Formulations

- 2.5× TB broth (125 grams of dehydrated TB culture medium/L, 10 ml glycerol/L)
- Gel filtration buffer (150 mM NaCl, 10 mM HEPES [pH 7.4])
- Lysis buffer (500 mM NaCl, 50 mM K_2_HPO_4_, 50 mM NaH_2_PO4), freshly supplemented with a Pierce^TM^ protease inhibitor cocktail tablet according to manufacturer’s instructions

### Kit

- Pierce™ BCA Protein Assay Kits (Cat. No. 23225, Thermo Scientific)

### Antibody for Western blotting

- BAX antibody (2D2) (Cat. No. SC-20067, Santa Cruz Biotechnology)
- Cytochrome *c* antibody (Cat. No. 54205-RBM6-P1, Thermo Scientific)
- m-IgGκ BP-HRP (Cat.No. sc-516102, Santa Cruz Biotechnology)

### Plasmids

- pTYB1-BAX [100 ng/μl], for expression of intein-tagged BAX (Suzuki et al., 2000)

### Materials for BAX validation assays (optional)

- (3-cholamidopropyl)dimethylammonio)-1-propanesulfonate (CHAPS) (Cat. No. C-080, Gold Biotechnology)
- 8-aminonapthalene-1,3,6-trisulfonic acid (ANTS) (Cat. No. A350, Thermo Fisher Scientific)
- BSA-fraction V (Cat. No. 12-660-9100GM, Fisher Scientific)
- Cardiolipin (18:1) (Cat. No. 710335C, Avanti Polar Lipids)
- Dodecylphosphocholine (DDPC) (Cat. No. 25629, Cayman Chemical)
- Brain phosphatidylserine (Porcine) (Cat. No. 840032C, Avanti Polar Lipids)
- Ethylenediaminetetraacetic acid (EDTA) (Cat. No. 798681-100G, Sigma-Aldrich)
- Ethylene glycol-bis(β-aminoethyl ether)-*N*,*N*,*N*′,*N*′-tetraacetic acid (EGTA) (Cat. No. E0396, Sigma-Aldrich)
- Egg phosphatidylcholine (Chicken) (Cat. No. 840051C, Avanti Polar Lipids)
- Egg phosphatidylethanoloamine (Chicken) (Cat. No. 840021C, Avanti Polar Lipids)
- Human BID-BH3, peptide (Cat. No. AS-61711)
- Human BIM-BH3, peptide IV (Cat. No. AS-62279)
- Liver phosphatidylinositol (Bovine) (Cat. No. 840042C, Avanti Polar Lipids)
- p-xylene-bis-pyridinium bromide (DPX) (Cat. No. X1525, Thermo Fisher)
- Recombinant Human BID (Caspase-8-cleaved) (Cat. No. 882-B8, R&D Systems)
- Sucrose (Cat. No. S0389, Sigma)
- Trehalose (Cat. No. 1673715, Sigma)

### Buffer Formulations for BAX validation assays (optional)

- Large unilamellar vesicles (LUVs) buffer (200 mM KCl, 5 mM MgCl_2_, 0.2 mM EDTA, 10 mM HEPES [pH 7.4])
- Trehalose isolation buffer (TIB) (200 mM trehalose, 68 mM sucrose, 10 mM HEPES pH 7.4, 10 mM KCl, 1 mM EDTA, 1 mM EGTA, 0.1% BSA-Fraction V), freshly supplemented with a Pierce^TM^ protease inhibitor cocktail tablet according to manufacturer’s instructions.

## Methods

### Reagents and buffer preparations

Prepare 100 mg/ml carbenicillin (1000×) stock (6 ml) by dissolving carbenicillin powder in distilled H_2_O (dH_2_O). Prepare 1 M IPTG (4 ml) stock by dissolving IPTG powder in dH2O. Store carbenicillin and IPTG stocks at −20°C and thaw as needed for immediate use.

Make 1× LB broth (1 L) by combining 25 g LB powder in 1 L of dH_2_O and thoroughly mix using a magnetic stir bar. Make 2.5× TB broth (4 L) by combining 500 g TB powder in 4 L dH2O, supplemented with 40 ml glycerol, and thoroughly mix using a magnetic stir bar. Sterilize both LB and TB stocks by autoclave and store at 4°C to prevent contamination. Prior to culturing bacteria, freshly supplement the LB and TB with carbenicillin to a final concentration of 100 μg/ml (1:1000-fold dilution).

Make 1× LB agar media (1 L) by combining 40 g of LB agar powder in 1 L dH_2_O and thoroughly mix with heat using a magnetic stir bar. Heat the agar mixture to boiling and maintain the boiling for 1 minute to completely dissolve the agar. Sterilize LB agar stock by autoclave and cool it down to 45–50°C. Supplement the LB agar stock with carbenicillin to a final concentration of 100 μg/ml and pour 20 ml into 10-cm petri dishes. Once the agar solidifies, store the LB agar plates at 4°C to prevent contamination.

Prepare lysis buffer (500 ml) by combining 83.3 ml 3M NaCl, 12.5 ml 2M K_2_HPO_4_ and 12.5 ml 2 M NaH_2_PO_4_ into 391.7 ml dH_2_O. Prepare gel filtration buffer (1 L) by combining 50 ml 3 M NaCl and 10 ml 1 M HEPES [pH 7.4] in 940 ml dH_2_O. CRITICAL: BAX can aggregate in the presence of detergent and common glassware can carry residual detergent from their cleaning. Best practice is to designate a set of glassware specifically for making lysis buffer and gel filtration buffer and avoid cleaning with detergents.

### Transform BL21 Star^TM^ (DE3) bacterial cells with pTYB1-BAX expression vector

Day 1. Thaw 12.5 μl BL21 cells on ice for 10 minutes before adding 50 ng of the construct, pTYB1-BAX. Gently flick the tube several times to mix and incubate on ice for 10 minutes. Heat shock the BL21 cells at 42°C for 45 seconds (using a water bath or heating block) and then immediately return the tube to ice for 2 minutes. Add 750 μl of LB broth to the tube and incubate the culture with agitation in a shaking incubator set to 220 rpm at 37°C for 45 minutes. (Note: for best agitation, have the tube positioned at an angle or perpendicular to the direction of the shaker). While the bacteria are incubating, place a prepared LB agar plate supplemented with 100 µg/mL carbenicillin in a 37°C incubator to warm the plate and eliminate condensation. Pellet the cells by centrifuging at 1,500 ×*g* for 2 minutes in a table-top centrifuge. Remove 700 μl of the supernatant using a pipet and resuspend the pellet in the remaining 50 μl of media. Pipet the BL21 cell suspension onto the prewarmed LB agar plate and spread the suspension evenly throughout the plate (using a sterile glass spreader, glass beads, pipet tip, or similar). Incubate the LB agar plate at 37°C for 14–16 hours. Note: this incubation step is commonly performed overnight and serves as a break point in the protocol.

### Expand BL21 (DE3) culture and induce expression of recombinant BAX

Day 2. Remove the culture plate from the incubator and inspect for colony growth. Keep the plate at 25°C until ready to proceed with the next step. Pick a single bacterial colony from the LB agar plate and inoculate into 400 ml of 1× LB freshly supplemented with 100 µg/ml carbenicillin in a 2 L flask. Expand this starter culture in a shaking incubator set to 225 rpm at 30°C for 14–16 hours. Note: typically, the plate is removed from the incubator in the morning and a colony is picked several hours later so that the incubation of the starter culture occurs overnight and aligns with the day schedule.

Day 3. In the morning, take 1 ml of bacterial culture to measure the optical density at 600 nm (OD_600_) using a spectrophotometer with 1 ml of dH2O as the blank solution. Continue to grow the starter culture until the OD_600_ value is approximately 1.5. Supplement the 400 ml of bacterial culture with 800 ml of 2.5× TB broth and 800 mL distilled H_2_O (dH_2_O), split evenly into two 2 L flasks, and incubate at 37°C shaking at 225 rpm to further expand the culture. For increased recombinant protein yield, 2–4 liters of TB culture is advised. Measure the OD_600_ each hour to monitor the population growth by removing 200 μl of the culture, diluting it with 800 μl dH_2_O in a cuvette (a 5-fold dilution). Use 1 ml of dH_2_O as a blank solution and adjust the OD_600_ value by the dilution factor (i.e., multiply by 5). Grow the culture to a target OD_600_ value of 2–3, which typically is achieved in 2–3 hours. Note: it is strongly recommended that investigators reserve a 100 μl aliquot of the confluent culture as a negative control before IPTG-induction to assess quality of BAX expression (“IPTG (−)”, see Figure 1A).

**Figure 1.**
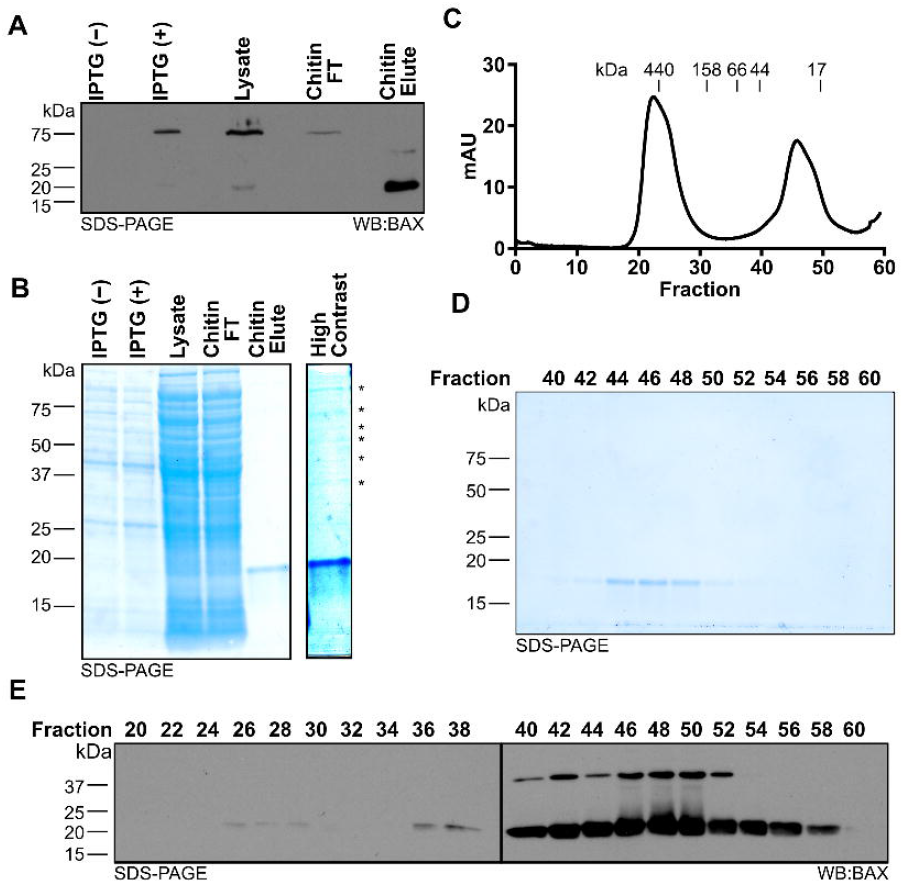
Systematic evaluation of BAX protein expression and purification. (A) Western blot detection of BAX for the samples collected throughout protein generation, which reveals intein-BAX expression induced by IPTG, intein-BAX captured by chitin column and BAX release by DTT-induced intein cleavage. (B) Left: protein staining of the samples collected throughout protein generation as in **A**; right: high contrast of chitin elute, which displays the presence of high molecular weight proteins noted with asterisk. (C) 280 nm absorbance values of sample exiting the HiLoad 16/600 Superdex 200 pg column. Molecular weights were determined by gel filtration chromatography standard. (D) Protein staining of fractions from **C**. (E) Western blot detection of BAX for the fractions from **C**.

Once the culture is at the target OD_600_, induce the expression of recombinant BAX by supplementing 1 mM IPTG (e.g., 1 ml of 1 M IPTG per liter of TB culture). Continue to incubate the bacterial culture in a shaking incubator set to 175 rpm at 30°C for 6 hours. Note: it is strongly recommended that investigators reserve a 100 μl aliquot of the confluent culture after IPTG-induction to assess quality of BAX expression (“IPTG (+)”, see Figure 1A).

### Pellet and lyse BL21 (DE3) cells

Pellet the BL21 (DE3) cells in 1-liter canisters by centrifuging at 4,000 ×*g*, at 4°C for 15 minutes in a high-speed centrifuge. Decant the supernatant and wash the bacterial pellet in the canister by resuspending with 25 ml of cold 1× PBS. Transfer the cell suspension to a 50 ml centrifuge tube and pellet the cells by centrifuging at 4,000 ×*g*, at 4°C for 45 minutes. Decant the PBS after centrifugation. Note: The cell pellet may be flash frozen in liquid nitrogen for future lysis. However, this may lead to reduced protein yield due to the additional freeze-thaw cycle.

Resuspend the BL21 cell pellet in 35 ml lysis buffer per 5 ml of pellet volume, and ensure cells are thoroughly resuspended by vigorous vortexing for optimal lysis efficiency. Combine the suspensions into a beaker placed on ice and continue to mix for 15 minutes using a magnetic stir bar. Freshly supplement the cell suspension with Pierce™ protease inhibitor cocktail following the manufacturer’s instruction. Keeping the beaker in ice, use a probe sonicator to lyse the cells using eight (8) cycles of the following program: run time at 2 minutes; PULSAR on at 15 seconds and PULSAR off at 2 minutes per cycle. Note: once the cells are lysed, it is strongly recommended that investigators reserve a 100 μl aliquot of cell lysate to evaluate the binding efficiency of recombinant BAX protein to the chitin resin (“Lysate”, see Figure 1A, third lane).

### Capture the recombinant BAX using a chitin affinity column

Recombinant BAX expressed by the pTYB1 vector is tagged with an intein domain for affinity chromatography using chitin beads. Thiol-induced self-cleavage of intein releases BAX from the beads for elution from the capture column.

Transfer the cell lysate into 40 ml Oak Ridge tubes and centrifuge in the high-speed centrifuge at 42,000 ×*g*, at 4°C for 1 hour to pellet cell debris. During the centrifugation, prepare the chitin column for capturing BAX. Gently swirl the chitin resin in its container to resuspend it. For each liter of TB culture, add 20 ml of the chitin resin slurry (equivalent to 10 ml of chitin resin bed volume) into a gravity column and allow the resin settle. Equilibrate the chitin column by washing twice with three (3) bed volumes of dH_2_O followed by washing twice with three (3) bed volumes of lysis buffer. Seal the chitin resin column when the lysis buffer level is approximately 0.5 cm higher than the resin layer.

Once the centrifugation is finished, load the supernatant directly onto the chitin column with a serological pipet, then unseal the chitin column to allow the supernatant to flow through the chitin resin. Collect the flow-through until the supernatant level is approximately 0.5 cm higher than the resin layer and then cap the column. Reapply the collected flow-through back onto the column and pass through again for additional BAX capture. Repeat once again for a total of three (3) passes of the supernatant through the column, capping after the final pass. Note: after the final capture, it is strongly recommended that investigators reserve a 100 μl aliquot of flow-through (FT) to evaluate the binding efficiency of recombinant BAX protein to the chitin resin (“Chitin FT”, see Figure 1A).

Wash the chitin column twice with three (3) bed volumes of lysis buffer, followed by equilibrating the column by washing three (3) times with 15 ml lysis Buffer freshly supplemented with 50 mM DTT. During the final wash, seal the chitin column when the lysis buffer level is approximately 0.5 cm higher than the resin layer. Incubate the chitin resin at 4°C for a minimum of 16 hours for on-column, DTT-induced cleavage of the intein tag and release of BAX from the chitin resin. Note: this step is commonly performed overnight and serves as a break point in the protocol.

At the end of day 3 during the on-column cleavage, equilibrate the HiLoad 16/600 Superdex 200 column with one (1) round of washing using one (1) column volume (120 ml) of dH2O, followed by an additional wash with one (1) column volume of gel filtration Buffer.

### Purify, concentrate, and store recombinant BAX

Day 4. Load 20 ml of lysis buffer onto the chitin column and pipet the chitin resin up and down several times to unpack the beads. Unseal the chitin column and collect the entire eluate (approximately 22–23 ml). Note: it is strongly recommended that investigators reserve a 100 μl aliquot of eluate to evaluate the efficacy of on-column cleavage (“Chitin Elute”, see Figure 1A). To further purify and isolate monomeric BAX, load an appropriate volume of the eluate onto the equilibrated HiLoad 16/600 Superdex 200 column according to the manufacturer’s instructions. The ÄKTA pure™ 25 system parameters are set as follows to conduct the chromatography: sample volume = 5 ml; system flow rate = 1 ml/min; fraction = 2 ml; alarm system pressure = 0.3 MPa. Perform the chromatography as many times as necessary to fully utilize the entire eluate.

To identify the BAX-containing fractions and determine purification efficiency, take 40 μl aliquot from alternating fractions, add Laemmli sample buffer, and subject to gel electrophoresis (i.e., SDS-PAGE). Stain protein gels with GelCode™ Blue Stain Reagent (or Coomassie equivalent) according to the manufacturer’s instructions to visualize protein bands (see Figure 1D). Pool the fractions containing only monomeric BAX and combine with fractions from each round of chromatography. Concentrate to a volume of approximately 3 ml using an Amicon^®^ Ultra-4 Centrifugal Filter according to the manufacturer’s instruction. Thoroughly mix the resulting 3 ml BAX sample with a pipet and quantify the BAX concentration using the Pierce™ BCA Protein Assay Kit. Based on the protein concentration and sample size volume, calculate the total yield of recombinant BAX and continue to concentrate BAX to an ideal concentration range of 30–40 μM with the same Amicon^®^ Ultra-4 Centrifugal Filter (Note: BAX may bind to the filter membrane, and using the same filter may avoid further loss of BAX protein). Aliquot the recombinant BAX into single-use aliquots (e.g., 20 μl per tube or suitable volume for the specific downstream applications), flash freeze using liquid nitrogen, and store at −80°C for future experimental interrogations.

### Large Unilamellar Vesicles (LUVs) permeabilization assays (optional)

Large unilamellar vesicles (LUVs) are prepared as previously described (Asciolla et al., 2012, Kuwana et al., 2002). Briefly, phosphatidylcholine, phosphatidylethanoloamine, phosphatidylserine, phosphatidylinositol and cardiolipin are combined at a ratio of 47:28:9:9:7 (5 mg total, for 7% cardiolipin LUVs) or 48:28:10:10:4 (5 mg total, for 4% cardiolipin LUVs), dried under nitrogen gas, and resuspended in LUV buffer containing a polyanionic dye (12.5 mM ANTS: 8-aminonaphthalene-1,3,6-trisulfonic acid) and cationic quencher (45 mM DPX: *p*-xylene-bis-pyridinium bromide) using a water bath sonicator. Unilamellar vesicles are formed by extrusion of the suspension through a 1.0 μm polycarbonate membrane. The unincorporated DPX and ANTS are removed by using a 10 ml Sepharose CL-2B gravity flow column. LUV preparations are only used for <2 weeks from when they are generated to avoid significant liposome degradation. For LUV permeabilization assays, using a 96 well format and 100 μl total volume per condition, BAX, BIM^BH3^, C8-BID and buffers are combined as indicated and analyzed for fluorescence (Excitation wavelength: 355 nm; Emission wavelength: 520 nm) using a multi-mode microplate reader. The percentage of release is calculated between the baseline provided by the buffer control and 100% release obtained by LUVs solubilized in 1% CHAPS.

### Mitochondrial Outer Membrane Permeabilization (MOMP) Assays (optional)

Mitochondria isolation and MOMP assays are performed as previously described (Renault et al., 2013). In brief, for the MOMP assays, 50 μg of mitochondria are treated with BAX, C8-BID, BIM^BH3^ peptide and buffers as indicated, and incubated in TIB supplemented with 100 mM KCl at 30°C for 1 hour. Every assay contains a 0.1% Triton X-100 control to determine maximal cytochrome *c* release. Samples are then centrifuged at 5,500 ×*g* for 10 minutes to separate into pellets and supernatants. The resulting pellet and supernatant fractions are analyzed SDS-PAGE and western blot to assess cytochrome *c* release.

### Fluorescence polarization ligand assay for monitoring BAX early activation (FLAMBE) assays (optional)

FLAMBE assays are performed as previously described (Gelles et al., 2022, Mohammed et al., 2022). Briefly, using a 96 well format and 100 μl total volume per condition, BAX, BIM^BH3^, C8-BID, buffers, and fluorescently-labelled BAK BH3 peptide (BAK^TAMRA^) are combined as indicated and analyzed for fluorescence (Excitation wavelength: 530 nm; Emission wavelength: 590 nm) using a multi-mode microplate reader fit with a polarization filter.

## Anticipated Results

### Systematic evaluation of BAX protein expression and purification

During this workflow, investigators should systematically collect protein samples at each key step to confirm protein expression, capture, and purification of monomeric BAX (highlighted in the Methods section). When using the pTYB1-BAX construct, the IPTG-induced intein-tagged BAX protein band is detected at 75 kDa (Figure 1A, first two lanes). Following three rounds of passing the bacterial lysate through the chitin column, the chitin column effectively captures the majority of recombinant BAX (Figure 1A, third and fourth lane). Investigators also need to confirm the release of BAX from the chitin column following the DTT-induced intein cleavage by sampling the eluate. As anticipated, the intein was removed during the DTT-induced cleavage process, and BAX from the chitin column elute was detected at 21 kDa (Figure 1A, last lane). The elute from chitin column is not entirely devoid of contaminating species as multiple higher molecular weight species were revealed by protein staining (Figure 1B, last lane), and so investigators need to perform additional purification using size exclusion chromatography.

The size exclusion chromatography technique separates contaminating species and other forms of BAX from monomeric BAX, allowing purify BAX in monomeric form. Prior to running the chromatography, we suggest investigators determine the resolution of their columns based on the manufacturer’s manual or run molecular standards. Following size exclusion chromatography, the chitin column elute should be fractioned into 2 distinct peaks. The first peak contains higher molecular weight species, and the second peak is expected to be a BAX monomer peak (Figure 1C). Then, investigators need to identify fractions containing monomeric BAX within the second peak and evaluate their purity by performing protein staining on alternating fractions from the size exclusion chromatography. Due to its relatively small size (21 kDa), BAX is typically eluted in later fractions (e.g., fraction 44–52, Figure 1D); however, the specific fractions containing BAX vary depending on the type of chromatography column and collection volume.

In the initial protein purification, investigators need to perform a western blot using BAX antibody to confirm that the observed protein is BAX. BAX bands should be detected at the same size as observed in the protein staining analysis (Figure 1E). It is worth nothing that BAX is able to dimerize during SDS-PAGE in a denaturing condition, so an additional BAX dimer band may be detected by western blot. However, the dimer band does not suggest the presence of BAX dimer in these fractions (e.g., 44–52) as the molecular standard suggests (Figure 1C). Since western blot has a far better detection limit, it is within expectation that BAX can be detected in early fractions (e.g., fraction 24–28), suggesting the presence of BAX higher molecular weight species in the chitin column elute. Collectively, fractions 44–52 were confirmed to exclusively contain monomeric BAX and pooled for concentrating and storage. Fractions 54–56 also exclusively contain monomeric BAX, but the total amount together is too little to improve the yield. Moreover, pooling fractions 54–56 for concentrating significantly prolongs the amount time spent on concentrating BAX, so we recommend not collect fractions 54–56. Typically, 2 liters of bacterial culture generate 2 ml of 30 μM (∼1.4 mg) recombinant BAX.

### Evaluate the functionality of recombinant BAX with Large Unilamellar Vesicles (LUVs) permeabilization assay

The apoptotic function of BAX is to form proteolipid pores on the OMM and induce MOMP. To evaluate the function of recombinant BAX, investigators need to conduct a series of membrane permeabilization assays using model membranes and/or isolated mitochondria. Large Unilamellar Vesicles (LUVs) are biochemically-defined liposomes that mimic the composition of the OMM and are well-suited for testing BAX function (Kuwana et al., 2002). Here, we present a series of different setting for investigators to evaluate the functionality of BAX using LUVs permeabilization assays. The first experiment is to test a range of BAX concentrations (e.g., 100–500 nM) for LUV permeabilization (Figure 2A). This experiment serves two purposes: 1) BAX protein exhibits concentration-dependent auto-activation and permeabilization should be observed at higher concentrations; 2) researchers can identify a suitable non-activating concentration of BAX for subsequent experiments. The BAX titration experiment will demonstrate auto-activation but not necessarily activation of the entire BAX population or the maximal amount of LUVs permeabilization for that concentration of BAX. Therefore, investigators can choose either BCL-2 direct activators (e.g., BID, BIM) for BAX or detergent to stimulate BAX for activation and assess the maximal BAX-mediated LUVs permeabilization.

**Figure 2.**
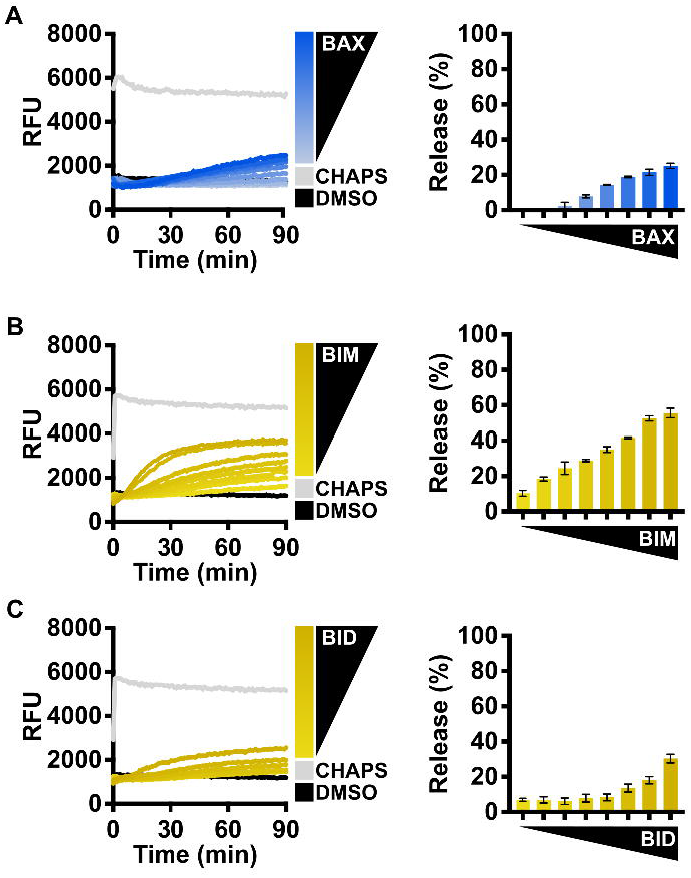
Evaluate BAX function with BCL-2 family direct activators. (A) Left: kinetics of LUVs permeabilized by concentration-dependent auto-activated BAX [105– 500 nM]; right: normalized endpoint of LUV permeabilization kinetics. (B) Left: kinetics of LUVs permeabilized by BAX [160 nM] with the presence of BIM^BH3^ [1:0–1:25 molar excess], demonstrating recombinant BAX is activated by BIM; right: normalized endpoint of LUV permeabilization. (C) Left: kinetics of LUV permeabilization by BAX [160 nM] with the presence of BID^BH3^ [1:0–1:75 molar excess], demonstrating recombinant BAX is activated by BID; right: normalized endpoint of LUV permeabilization. Kinetic data are represented as mean of triplicates; error bars denote S.D.

### BCL-2 direct activator-mediated BAX activation

The activity of BAX is primarily regulated by its interaction with other BCL-2 family proteins, and so we recommend investigators conduct LUVs permeabilization assays with different forms of BCL-2 direct activators for BAX. In cells, BIM is the predominant activator of BAX (Sarosiek et al., 2013), and so we suggest investigators first test BAX treated with a titration of BIM^BH3^ peptide (Figure 2B). This experiment serves two purposes: 1) BIM^BH3^ peptide exhibits a concentration-dependent activation on BAX membrane permeabilization function, confirming BAX responds to BH3 stimulation; 2) investigators can identify a suitable molecular ratio for future experiments (e.g., synergistic/antagonistic test with BAX activity modulator).

Another direct activator, BID, typically activates BAX following death receptor signaling (Li et al., 1998), and so investigators may choose BID^BH3^ peptide if their primary interest is to study BID-mediated BAX activation, albeit it requires much higher concentration compared to BIM^BH3^ peptide (Figure 2C). The potency of BH3 peptide largely depends on the length of peptide (Gelles et al., 2022). The commercially available BID^BH3^ peptide is 6 amino acids shorter than the BIM^BH3^ peptide, and so we anticipate BID^BH3^ to show decreased potency to activate BAX.

The caspase-8 cleaved BID (C8-BID) protein has been historically utilized as the major *in vitro* direct activator for BAX and so investigators may alternatively utilize C8-BID to study BAX activation. C8-BID has increased potency in activating BAX at nanomolar range due to its tertiary structure (Figure 3A). Several studies suggest that cardiolipin recruits C8-BID to OMM and promotes the function of tBID to activate BAX (Lutter et al., 2000, Kuwana et al., 2002), and so it is anticipated that C8-BID performs better at activating BAX to permeabilize higher amount of membrane at increased rate when LUVs contain a higher percentage of cardiolipin (i.e., 7%) (Figure 3A–C). For investigators primarily using BIM to study BAX activation, there is no notable difference between using LUVs with 7% cardiolipin and 4% cardiolipin (Figure 3D).

**Figure 3.**
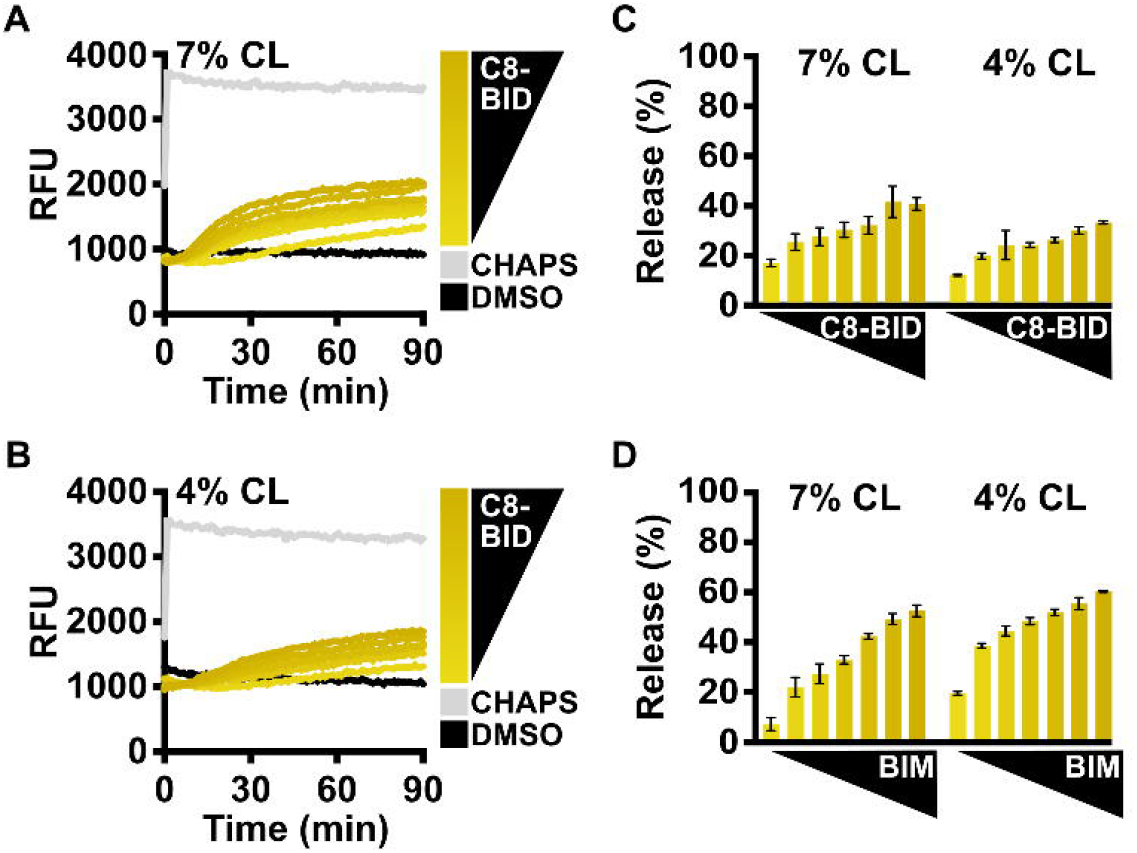
Compare BCL-2 direct activators using LUVs with different cardiolipin percentages. (A) Kinetics of LUVs permeabilized by BAX [160 nM] with the presence of C8-BID [33–250 nM] using the 7% cardiolipin (“CL”) LUVs, demonstrating recombinant BAX is activated by C8-BID. (B) Kinetics of LUVs permeabilized by BAX [160 nM] with the presence of C8-BID [33–250 nM] using the 4% cardiolipin (“CL”) LUVs. (C) Normalized endpoint of LUV permeabilization from **A** and **B**. (D) Normalized endpoint of LUVs permeabilized by BAX [160 nM] activated by BIM^BH3^ [1:0–1:25 molar excess] using 7% or 4% cardiolipin (“CL”) LUVs. Kinetic data are represented as mean of triplicates; error bars denote S.D.

#### Detergent-mediated BAX activation

Investigators can also use *n*-dodecyl-phosphocholine (DDPC), a detergent that has been reported to create homogenous BAX oligomers mimicking physiological activation (Hauseman et al., 2020), to assess the enhanced LUV permeabilization for several concentrations of BAX (Figure 4A). Alternatively, the detergent octyl-β-glucoside (OG) has been long-established to activate BAX (Hsu and Youle, 1998) and can be utilized for assessing real-time BAX activation and LUV permeabilization at several concentrations of BAX (Figure 4B). Collectively, the aim of performing these assays is to demonstrate that the BAX protein is functional and to determine an appropriate concentration of BAX that will not auto-activate but will robustly permeabilize LUVs and exhibit an appropriate dynamic range for subsequent experiments.

**Figure 4.**
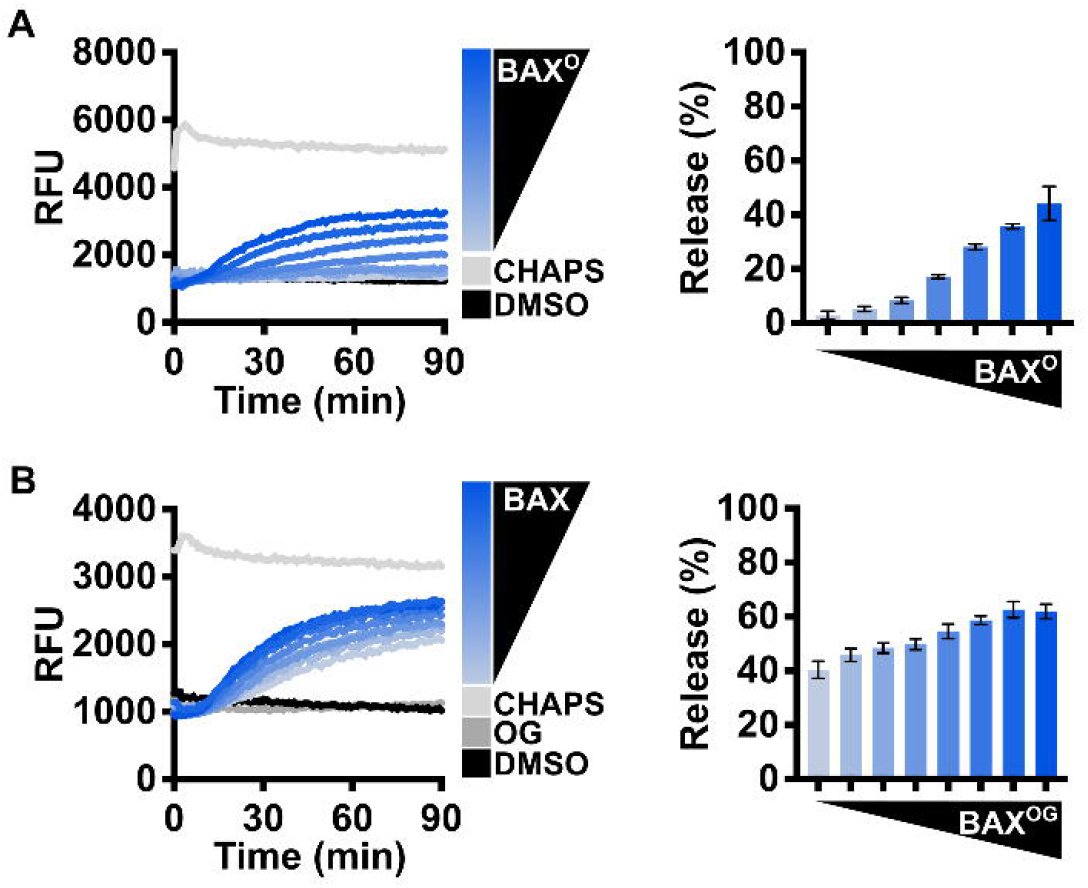
Evaluate BAX functionality with non-physiological BAX activator-detergents. (A) Left: kinetics of LUVs permeabilized by DDPC-induced BAX oligomers (“BAX^O^”, [44–500 nM]); right: normalized endpoint of LUVs permeabilized by BAX^O^. BAX^O^ was produced by incubating BAX with DDPC [1 mM] for 14–16 hours at 4°C. (B) Left: kinetics of LUVs permeabilized by OG-activated BAX [OG: 0.2%; BAX: 105–500 nM]; right: normalized endpoint of LUVs permeabilized by OG-activated BAX. Kinetic data are represented as mean of triplicates; error bars denote S.D.

Collectively, these results are fairly reproducible and so if a batch of BAX protein deviates from the expected results, it may suggest issues with protein quality, purity, or concentration.

### Evaluate BAX functionality with isolated mitochondria

In cells, following BAX-mediated MOMP, cytochrome *c* is released into the cytosol to trigger caspase activation. Thus, this field has been using isolated mitochondria as a fundamental standard to assess cytochrome *c* release for studying BCL-2 family protein function. Mitochondria should be freshly isolated from *BAX^−/−^ BAK^−/−^* MEFs and subject to MOMP on the same day to avoid cytochrome *c* leakage. Similar to LUVs permeabilization assays, investigators first need to test a range of BAX concentrations [10–50 nM], and BAX should active in a concentration-dependent manner to permeabilize mitochondria and result in cytochrome *c* release into supernatant. This experiment serves to identify a suitable BAX concentration for subsequent BAX activation study (Figure 4A). Investigators then need to use the identified BAX concentration to recapitulate synergy between BAX and BIM^BH3^ peptide or C8-BID protein to establish appropriate molecular ratio for their future research (Figure 5A).

**Figure 5.**
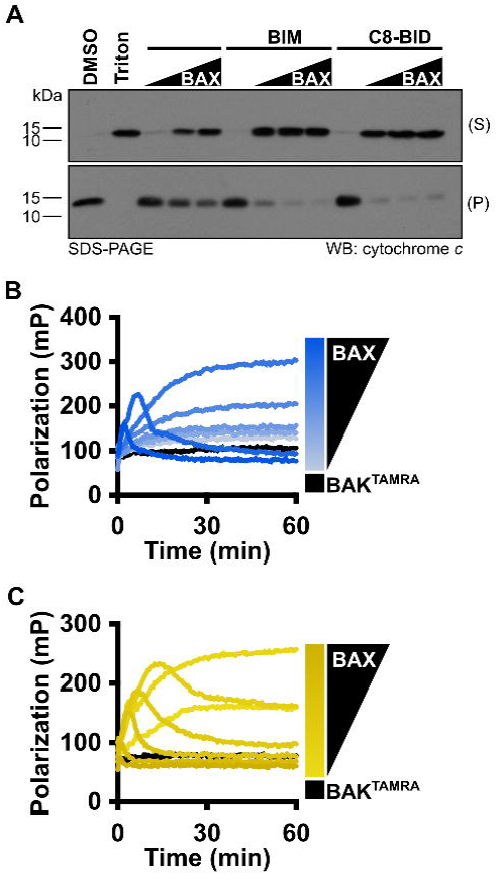
MOMP and FLAMBE assays are valuable techniques to study BAX activation. (A) Isolated mitochondria were incubated with BAX [5, 25, 50 nM], C8-BID [25 nM] and BIM^BH3^ [2.5 μM] peptide as indicated. Mitochondria were centrifuged and the resulting supernatant (S) and pellet (P) were subject to western blot detection of cytochrome *c*. (B) BAK^TAMRA^ [50 nM] was added to BAX [17–200 nM] and fluorescence polarization was measured. (C) BAK^TAMRA^ [50 nM] was added to BAX [60 nM] in the presence of BIM^BH3^ [0.26–2 μM] and fluorescence polarization was measured. Kinetic data are represented as mean of replicates.

### Study BAX activation with fluorescent polarization assays

Before BAX is fully competent to induce membrane permeabilization, it must undergo a series of conformational change in the cytosol and OMM. For investigators who want to gain insight into the BAX early activation process, we recommend using the FLAMBE assay (Mohammed et al., 2022). Using fluorescently labelled BAK^BH3^ peptide capable of binding to the BAX BC groove, FLAMBE measures activation-induced release of BAK^BH3^ peptide to monitor BAX intramolecular conformational change in the BAX early activation. It can detect both concentration-dependent BAX auto-activation and BH3-stimulated BAX activation indicated by decrease in polarization signals (Figure 5B). Collectively, these assays study BAX activation from different perspective and corroborate with LUVs permeabilization assay to comprehend BAX apoptotic function research.

### BAX handling and storage

Given BAX activity, protein stocks have a tendency to form multimers and aggregates during storage. Therefore, proper handling and storing of BAX stocks is key to preserving a functional and responsive BAX population for downstream investigations and consistent experimental results. We recommend concentrating BAX to a range of 30–40 μM since overconcentrating BAX can result in protein aggregation and subsequently a population that is less sensitive to BH3 stimulation (Figure 6A–B). Additionally, we observed BAX dimerization at high concentrations (data not shown), which may interfere with certain technique analyses (e.g., NMR and ITC). NMR and ITC analyses usually require higher protein concentrations to generate adequate signal and typically 40 μM stocks are sufficient for these techniques. To store BAX, we recommend investigators aliquot BAX into single use tubes (e.g., 10–20 μl for plate-based assays), flash freeze with liquid nitrogen, and store them at −80°C for future use. Investigators need to note that larger aliquots can slow the flash freeze, resulting in protein precipitation and loss. BAX stored in −80°C only exhibits a minor reduction in BAX membrane-permeabilization activity but maintains its response to BH3 stimulation (Figure 6C). In contrast, BAX stored at 4°C exhibits a notable loss in BAX function at the same concentration and a reduced response to BH3 stimulation (Figure 6D). In conclusion, we suggest concentrate BAX to a range of 30–40 μM, aliquot in small volume, flash freeze with liquid nitrogen, and store in −80°C for future use.

**Figure 6.**
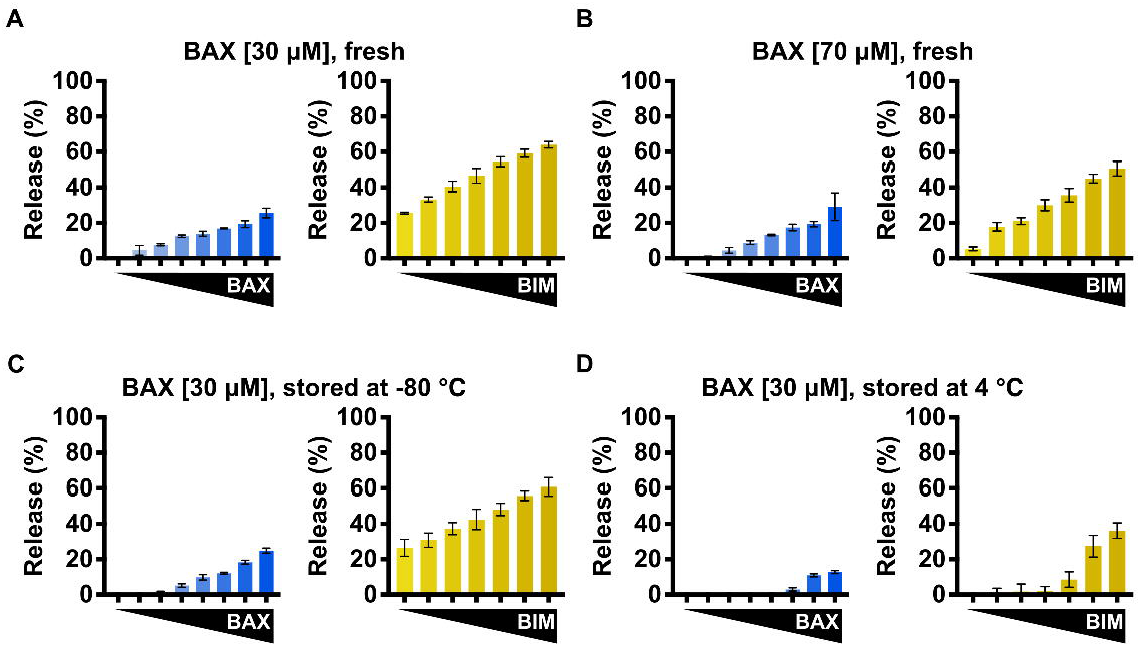
BAX storage conditions impact BAX functionality. (A–D) Left: normalized endpoint LUV permeabilization by auto-activated BAX [105–500 nM]; right: normalized endpoint LUV permeabilization by BIM^BH3^-activated BAX [160 nM] with the presence of BIM^BH3^ [1:0–1:25M excess]. (A) Purified BAX was concentrated to 30 μM and immediately subject to LUV assay. (B) Purified BAX was concentrated to 70 μM and immediately subject to LUV assay. (C) Purified BAX was concentrated to 30 μM and stored for 1 week at −80 °C before subject to LUV assay. (D) Purified BAX was concentrated to 30 μM and stored for 1 week at 4°C before subject to LUV assay. Error bars denote S.D.

## Discussion

The mitochondrial pathway of apoptosis is fundamental for tissue homeostasis, and dysregulated control of this pathway is implicated in various human diseases including cancer, heart failure, and neurogenerative disorders (Spitz and Gavathiotis, 2022). Given the critical role of BCL-2 family proteins in the regulation of apoptosis, research has been dedicated to developing small molecules to inhibit the anti-apoptotic BCL-2 proteins (i.e., BH3 mimetics). Recent mechanistic studies of BAX molecular regulation have identified several BAX regulatory regions, providing key structural insights that are aiding in the development of small molecules capable of directly targeting BAX and modulating its activity (Pritz et al., 2017, Garner et al., 2019, Gavathiotis et al., 2012). These efforts were built upon the ability to generate recombinant BAX for structure-function interrogations and small molecule screens.

Despite the growing number of publications utilizing recombinant BAX, Methods sections are not sufficiently informative to learn the technique *de novo* and there remains a lack of detailed and instructional protocols aimed at investigators who are not already familiar with generating BAX protein. In this work, we present a comprehensive protocol for expressing, purifying, and storing functional recombinant BAX protein with a yield of ∼1.4 mg per 2 liters of bacterial culture. To ensure the quality of each batch of BAX, we have recommend and provided examples of quality control and BAX activation assays. Additionally, given the high purity level of recombinant BAX, our two-step chromatography protocol can be easily adapted to generate N^15^-or C^13^-labelled BAX for HSQC NMR study.

Moreover, this protocol is suitable for purification of BAX mutants following mutagenesis of the expression vector. It is worth noting that some structural mutants of BAX are created by introducing cysteine residues to form disulfide bonds, and these BAX mutants must be oxidized by incubating in an oxidative environment to generate the disulfide tether (Gavathiotis et al., 2010, Czabotar et al., 2013, Gelles et al., 2022). While not being physiological activators, both DDPC and OG are beneficial for evaluating loss-of-function BAX mutants as they may bypass the certain structural requirements for BAX oligomerization and serve as robust positive controls for BAX activity.

Collectively, we optimized handling and storage condition for BAX, and BAX can be stored at −80°C for up to 6 months with consistent and reproducible function. We recommend a range of 30–40 μM for BAX storage to avoid BAX multimerization, and this range is suitable for most techniques essential for BAX research.

## Data availability statement

The original contributions presented in the study are included in the article and Supplementary Material, further inquiries can be directed to the corresponding author.

## Author contributions

Conceptualization, Y.C., J.D.G., and J.E.C.; methodology, Y.C., J.D.G., and J.N.M.; investigation, Y.C., J.D.G., and J.N.M.; writing, Y.C., J.D.G., and J.E.C.; resources, Y.C.; visualization, Y.C., J.D.G.; funding acquisition, J.E.C.; supervision: J.E.C.

## Acknowledgments

We would like to acknowledge the TCI Shared Resources and the Department of Oncological Sciences for research support.

